# Pre-treatment of Nile tilapia (*Oreochromis niloticus*) with ozone nanobubbles improve efficacy of heat-killed *Streptococcus agalactiae* immersion vaccine

**DOI:** 10.1101/2021.12.13.472363

**Authors:** Nguyen Vu Linh, Le Thanh Dien, Pattiya Sangpo, Saengchan Senapin, Anat Thapinta, Wattana Panphut, Sophie St-Hilaire, Channarong Rodkhum, Ha Thanh Dong

## Abstract

Nanobubble technology has shown appealing technical benefits and potential applications in aquaculture. We recently found that treatment with ozone nanobubbles (NB-O_3_) activated expression of several immune-related genes leading to effective response to subsequent exposure to fish pathogens. In this study, we investigated whether pre-treatment of Nile tilapia (*Oreochromis niloticus*) with NB-O_3_ can enhance specific immune responses and improve efficacy of immersion vaccination against *Streptococcus agalactiae*. Spleen and head kidney of fish in the vaccinated groups showed a substantial upregulation in expression levels of three immunoglobulin classes(*IgM*, *IgD*, and *IgT*) compared with the unvaccinated control groups. At day 21 post-immunization, the relative expression was greatest (approx. 3.2 to 4.1 folds). Both systemic and mucosal IgM antibodies were elicited in vaccinated groups. As the result, the cumulative survival rate of the vaccinated groups was found to be higher than that of the unvaccinated groups, with a relative percent survival (RPS) ranging from 52.9-70.5%. However, fish in the vaccinated groups that received pre-treatment with NB-O_3_, bacterial antigen uptakes, expression levels of *IgM*, *IgD*, and *IgT*, as well as the specific-IgM antibody levels and percent survival, were all slightly or significantly higher than that of the vaccinated group without pre-treatment with NB-O_3_. Taken together, our findings suggest that utilizing pre-treatment with NB-O_3_ may improve the immune response and efficacy of immersion vaccination in Nile tilapia.

**Highlights:**

- Immune response and efficacy of a heat-killed *Streptococcus agalactiae* immersion vaccine for Nile tilapia with and without pre-treatment with NB-O_3_ were accessed.
- Bacterial antigen uptake in the NB-O_3_-VAC compared to the AT-VAC groups was increased 1.32 and 1.80-fold at 3 and 6 h post-vaccination, respectively.
- Vaccinated group that received pre-treatment with NB-O_3_ had slightly to significantly higher levels of *IgM*, *IgD*, and *IgT* mRNA expression; IgM levels; and survival rate.
- Pre-treatment with NB-O_3_ may be a novel strategy for improving efficacy of immersion vaccine in aquaculture

## 1. Introduction

Aquaculture has grown at an unprecedented rate worldwide over the past two decades [1]. This expansion has resulted in larger and more numerous farms within watersheds, which enhances the risk of host-dependent pathogen transmission and makes disease management more challenging [2–4]. Disease outbreaks are a major cause of economic losses in aquaculture, including Nile tilapia (*Oreochromis niloticus*) [5–8]. As a result of these developments, the use of chemotherapy to control diseases has increased. Recently, numerous antibiotics have been used in aquaculture to minimize losses due to bacterial diseases. When an antibiotic treatment is unsuccessful, the farmer typically switches to another antibiotic or increases the dose of the medication, both of which result in greater antibiotic usage [9], which is problematic for the development of antimicrobial resistance (AMR).

Disease prevention is the most rational approach to resolving the problems associated with antibiotic treatments, as it reduces the need for these products. Reduced antibiotic usage in aquatic systems will eventually decrease the risk of AMR in these sectors, which may directly affect AMR risk in human populations. Prevention of disease can be accomplished through a number of ways, which include reducing exposure to pathogens and/or improving host resistance to disease [10,11]. The latter can be achieved through vaccination [12]. To date, vaccination has been shown to be the most efficient method for combating pathogenic infections or conferring struggle to target pathogens. Numerous successful vaccines have been produced that provide effective protection in fish, including subunit vaccines, inactivated vaccines, DNA vaccines, and vaccines with live attenuation [13–16]. However, issues of vaccination also limit their use. Historically, immunizations have been administered through injection, which is time consuming and difficult to deliver to young fish [17]. While oral and immersion vaccinations are simple to administer with minimum stress to the fish, they often generate limited immune responses [12,18–20]. Improving immersion vaccines would go a long way to preventing infectious diseases on fish farms [12].

A new technology that injects nanobubbles into liquids and helps reduce bacterial counts in water may reduce pathogen burdens on fish farms [21–24]. In our previous investigations, ozone nanobubbles were found to be efficient in reducing bacterial concentrations in water and upregulating the innate immune system of fish, resulting in increased fish survival during pathogenic infections. When Nile tilapia were infected with a pathogenic multidrug-resistant *Aeromonas hydrophila*, they displayed a higher survival rate when exposed to treatments with ozone nanobubbles (NB-O_3_) [25–27]. Given the stimulation of the innate immune system in the treated groups, we hypothesized that this technology might improve the efficacy of immersion vaccines. This study aimed to investigate: 1) whether treatment of Nile tilapia with NB-O_3_ can enhance specific immune responses to vaccine, and 2) whether simultaneous treatment with NB-O_3_ can improve the efficacy of immersion vaccination against *S. agalactiae*.

## 2. Materials and Methods

### 2.1 Animals and ethical issues

A total of 360 apparently healthy Nile tilapia fish were provided by a tilapia hatchery (Department of Fisheries, Thailand). Experimental fish were maintained in fiberglass tanks (100 L), which were continuously aerated for 2 weeks and equipped with a cotton filter before the vaccination trials. Prior to conducting additional experiments, 10 randomly selected fish were subjected to bacterial and parasite examinations to guarantee their health. The Thai Institutional Animal Care and Use Committee authorized all animal operations (approval no. MUSC64-024-573).

### 2.2 Bacterial culture and heat-killed vaccine preparation

*Streptococcus agalactiae* strain 2809, identified from a tilapia field outbreak, was used for this research (Centex Shrimp, Mahidol University, Thailand). It was retrieved from frozen glycerol stocks and cultivated for 24 h at 28 °C on tryptic soy agar (TSA, Becton, Dickinson and Company, USA), followed by culturing in 100 mL of tryptic soy broth (TSB, Becton, Dickinson and Company, USA) for 18 h. The bacterial cells were inactivated using the heat-killed method at 56 °C for 30 min in a water bath [28]. To confirm bacterial inactivation, an aliquot of 0.1 mL of killed bacterial suspension was plated onto TSA and *S. agalactiae* selective agar bases (SSA, HiMedia, India) and incubated for 48 h at 28 °C. The absence of bacterial growth indicated successful inactivation. The heat inactivated *S. agalactiae* without adjuvant was used as immersion vaccine in this study.

### 2.3 Fish vaccination and challenge

Two weeks after acclimation, 360 fish (15.62 ± 0.45 g) were randomly allocated into 4 experimental groups: ozone nanobubbles without vaccine (NB-O_3_-noVAC) group (G1), ozone nanobubbles with vaccine (NB-O_3_-VAC) group (G2), an air-stone with vaccine (AT-VAC) group (G3), and an air-stone without vaccine (AT-noVAC) group (G4). Each treatment was performed in duplicates with 45 fish per tank. Firstly, the nanobubble tanks (G1, G2) were subjected to NB-O_3_ for 10 min according to a previously reported protocol [27]. After 3 h, the vaccinated groups (G2, G3) were immunized with heat-killed *S. agalactiae* vaccine (1.56 × 10^9^ CFU/mL) by adding 1 L of inactivated vaccine to each tank containing 50 L of water and 45 fish to reach a final concentration of 1.67 x 10^7^ CFU/mL. The other non-vaccinated groups (G1, G4) were carried out in the same manner using TSB without inactivated bacteria as the control group. Following a 12-hour immersion vaccination, fish were transferred to new aeration tanks and maintained at 30 ± 1 °C for 21 d.

The efficiency of vaccines against *S. agalactiae* was assessed using an experimental challenge with *S. agalactiae* on day 21 after vaccination. The experimental trials were conducted in 100 L dechlorinated tap water tanks, including 20 fish per tank. The fish immunized with heat-killed vaccines (n = 20) were injected intraperitoneally with *S. agalactiae* at a dose of 10^7^ CFU/fish. By contrast, the NB-O_3_-noVAC and AT-noVAC groups received injections of 0.1 mL of 1× PBS. Fish mortality was monitored for 14 d (Fig. 1).

**Fig. 1.**
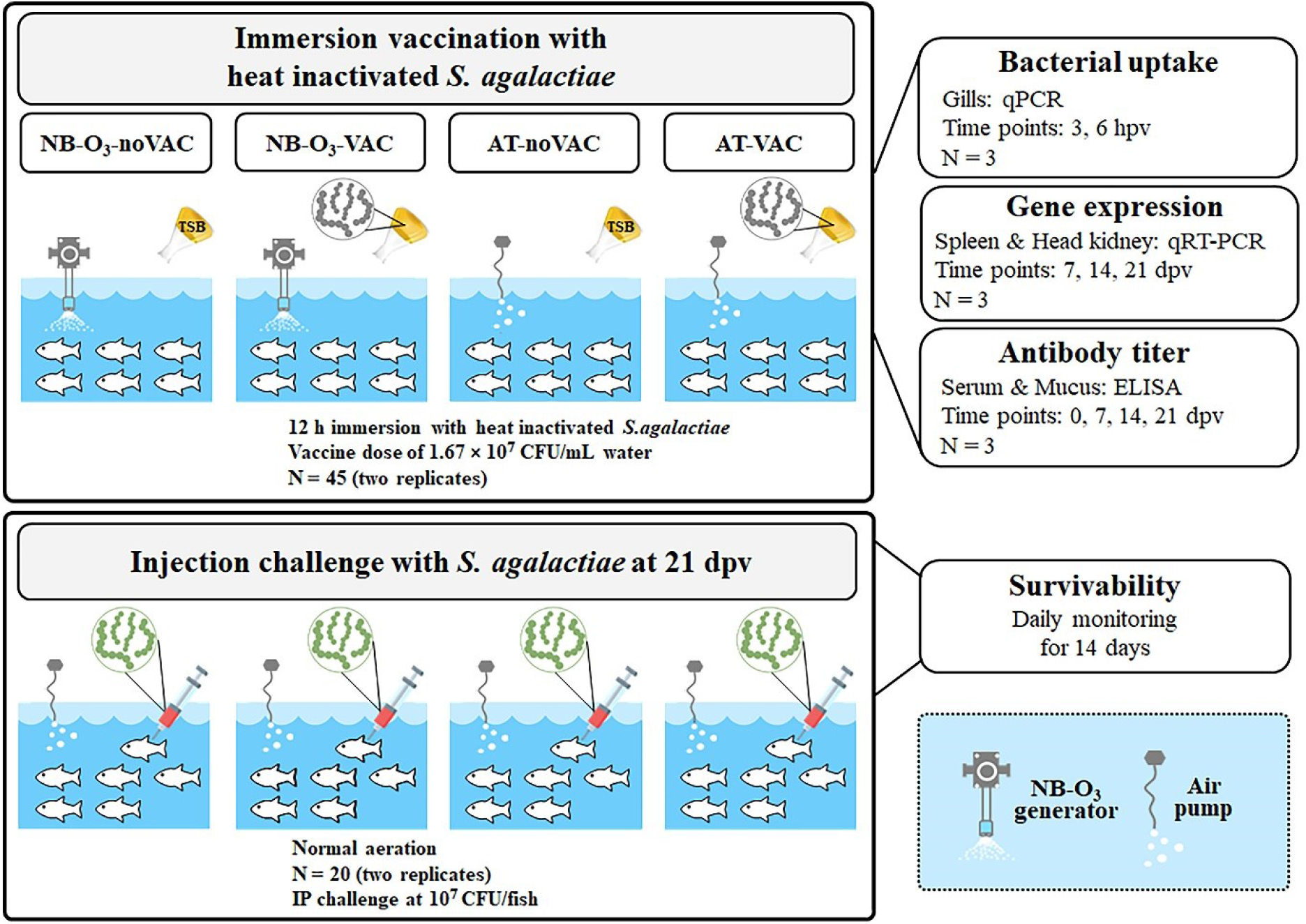
Schematic diagram of experimental design illustrating before and after immersion vaccination, and Nile tilapia species used. AT-noVAC, air-stone with no vaccine; NB-O_3_-noVAC, ozone nanobubbles without vaccine; AT-VAC, air-stone with vaccine; NB-O_3_-VAC, ozone nanobubbles with vaccine. TSB: Tryptic Soy Broth media.

### 2.4 Sample collection

To investigate the uptake of vaccine into fish bodies of Nile tilapia through gills, six representative fish (six biological replicates/group) were selected at different periods (3 and 6 h) post-immersion vaccination. Prior to sample collection, the fish were euthanized with a lethal dose of clove oil (250 ppm). The gill tissues from each fish were collected and stored in 95% ethanol at a ratio of 1:10 (v/v) tissue:ethanol until further analysis.

To collect the mucus and serum samples for enzyme-linked immunosorbent assay (ELISA), six random fish were selected at different intervals: days 0 (baseline), 7, 14, and 21. Mucus samples were collected by gently rubbing the fish in a plastic bag that contained 1 mL of 1× PBS and 0.02% sodium azide [29]. Blood samples were obtained from the tail vein of fish using a syringe fitted with a 23-G needle. The serum and mucus were then collected and kept at –20 °C for further analysis.

To analyze mRNA expression of three immunoglobulin genes encoding *IgM*, *IgD*, and *IgT*, the fish tissues (spleen and head kidney) were collected at different periods (days 7, 14, and 21) after immunization. Investigated tissues (40–50 mg) from six randomly chosen fish (mentioned earlier) were collected and stored in sterile tubes supplemented with 200 μL Trizol (Invitrogen, USA) at −20 °C until examinations.

### 2.5 qPCR assay for quantifying *S. agalactiae*

Quantitative PCR was performed according to the protocol reported by Leigh et al. [30]. Primers *SagroEL-F/R*, which targets the *groEL* gene of *S. agalactiae* (accession number EU003621). A 142-bp product amplified from *S. agalactiae* 2809 was cloned into pGEM and the recombinant plasmid, namely pSNB1, was used as a positive control and for standard curve construction. Serial dilutions of the pSNB1 plasmid spiked with 200 ng of tilapia DNA were used to construct a standard curve for quantifying *S. agalactiae*. Fish gill DNA was isolated using the conventional phenol-chloroform method [31,32] and 200 ng of each DNA sample was subjected to qPCR assays using the CFX Connect™ Real-time System (Bio-Rad, USA). The resulting C_q_ value was used to compute bacterial DNA in the fish gills using the equation: copy number = 10^(Ct – Intercept)/Slope^. qPCR for each template was performed in triplicate and calculated as bacterial load per 1 μg DNA template.

### 2.6 qPCR assay for immune gene expression study

Total RNA was isolated from tissue samples (spleen and head kidney) using the Trizol method according to the manufacturer’s procedures. First-strand complementary DNA synthesis and qPCR were performed following the procedures reported by Linh et al. [27]. The primers specific for tilapia *IgM, IgD, IgT*, and *β-actin* used for qPCR are listed in Table 1. The 2^-ΔΔCt^ method was used to analyze relative gene expression data [33]. Transcript levels of AT-noVAC groups on day 7 were set at 1.

**Table 1:**
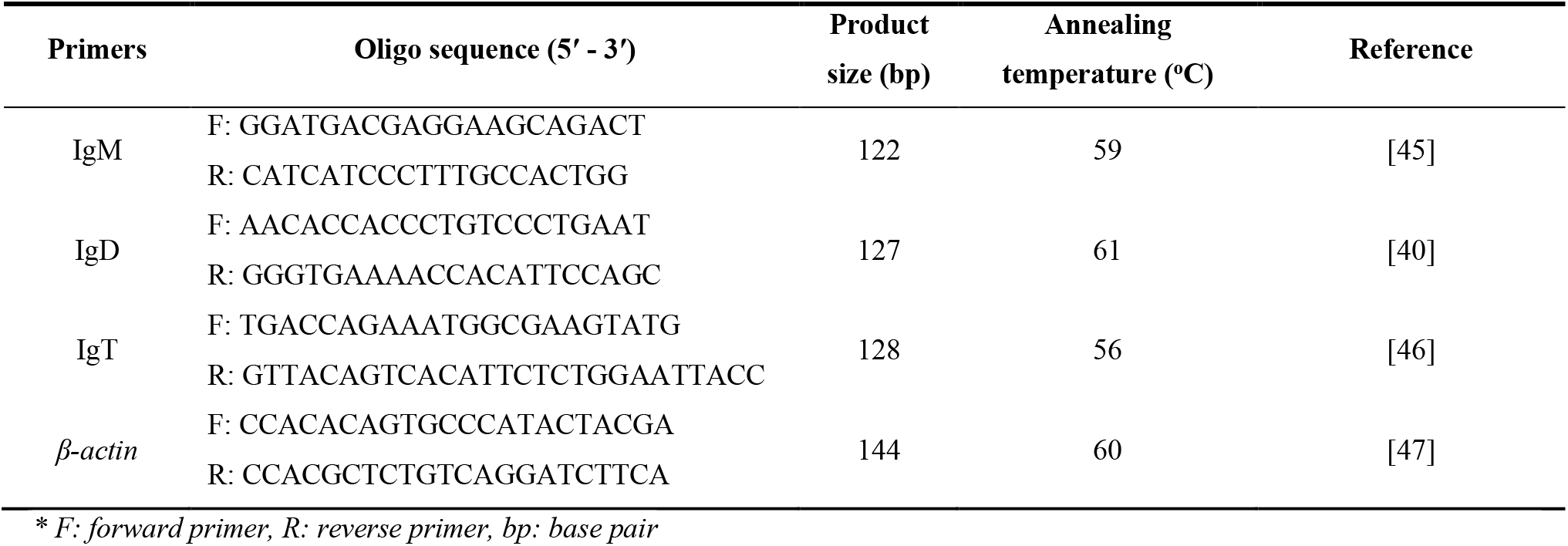
Primers used in this study.

### 2.7 Serum and mucus antibody assays

The mucosal and systemic IgM antibody levels of Nile tilapia were detected using ELISA. Mucus and serum ELISA assays were performed in the same manner as previously reported [27], with minor modifications. Briefly, mucus or serum samples were collected from six representative fish of each time point (days 0, 7, 14, and 21). Two-fold serial dilutions were performed to determine the optimal dilution. The ELISA dilutions for mucus and serum were 1:16 and 1:512, respectively.

### 2.8 Statistical analysis

SPSS program (ver. 22.0) was used to conduct all statistical analyses. The Kaplan–Meier method was used to evaluate the survival rates in challenge trials, and a log-rank test was used to compare the treatment groups. One-way analysis of variance (ANOVA) was used to evaluate expression of immunoglobulin genes. Duncan’s post hoc tests were used to compare mean values. The Kruskal– Wallis test was used to examine the ELISA data. Bonferroni test was used to compare various groups. *P* ≤ 0.05 was considered statistically significant.

## 3. Results

### 3.1 Quantification of bacterial uptake into the fish gills after immunization

The qPCR for *S. agalactiae* performed in the current research showed a detection limit of 100 copies/μL of target template with an amplification efficiency of 90.2% and an R^2^ = 0.991. The mean C_q_ ± standard deviation (SD) for the detection limit was 36.64 ± 0.54 (Fig. 2). That is to say, samples with C_q_ ≤ 36.14 were considered *S. agalactiae* positive. The mean bacterial uptake ± SD in the NB-O_3_-VAC groups was 2032.40 ± 2053.45 and 5669.50 ± 2763.31 per 1 μg DNA template, whereas a lower value (1539.61 ± 585.91 and 3223.92 ± 970.96) was observed in the AT-VAC groups at 3 and 6 h post-immunization. The bacterial DNA present in the gills in the NB-O_3_-VAC was 1.32 and 1.80-fold higher compared to that in the AT-VAC groups after 3 and 6 h of vaccination, respectively (Table 2).

**Fig. 2.**
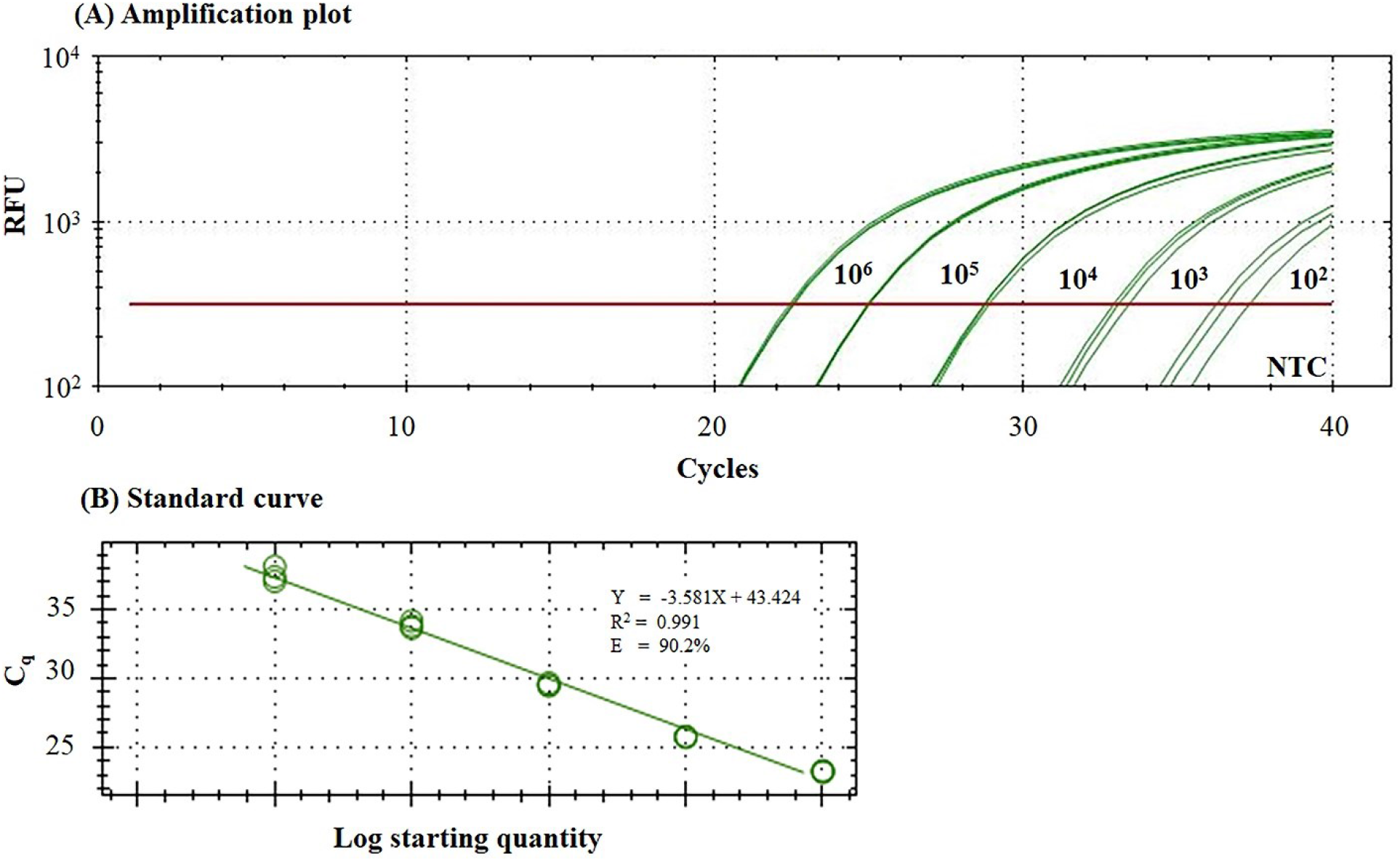
Detection of *Streptococcus agalactiae* using quantitative polymerase chain reaction (qPCR). (A) Amplification plots of positive control plasmid pSNB1 serial dilutions from 10^6^ to 10^2^ copies with 200 ng spiked fish DNA in each reaction. Three technical duplicates are included for each dilution. (B) A standard curve is constructed by plotting C_q_ values versus log_10_ concentrations. Formula for calculating copy number, R^2^, and E value are presented in the box.

**Table 2:** Quantification of *Streptococcus agalactiae* DNA on gill tissues by quantitative polymerase chain reaction (qPCR)

| Samples | 3 h post-vaccination |  | 6 h post-vaccination |  |
| --- | --- | --- | --- | --- |
|  | Mean C<sub>q</sub> | Bacterial loads/1 µg DNA template | Mean C<sub>q</sub> | Bacterial loads/1 µg DNA template |
| **NB-O<sub>3</sub> and vaccine groups** |  |  |  |  |
| NB-O<sub>3</sub>-VAC-1 | 32.55 | 5445.77 | 32.07 | 7419.79 |
| NB-O<sub>3</sub>-VAC-2 | 36.74 | - | 33.83 | 2393.43 |
| NB-O<sub>3</sub>-VAC-3 | 35.82 | 664.35 | 31.56 | 10270.22 |
| NB-O<sub>3</sub>-VAC-4 | 35.82 | 664.35 | 32.82 | 4550.15 |
| NB-O<sub>3</sub>-VAC-5 | 35.34 | 905.39 | 32.83 | 4531.08 |
| NB-O<sub>3</sub>-VAC-6 | 33.77 | 2482.13 | 32.73 | 4852.34 |
| Mean |  | 2032.40 ± 2053.45 |  | 5669.50 ± 2763.31 |
| **AT and vaccine groups** |  |  |  |  |
| AT-VAC-1 | 37.13 | - | 33.20 | 3570.26 |
| AT-VAC-2 | 36.26 | - | 33.21 | 3559.03 |
| AT-VAC-3 | 34.30 | 1765.36 | 33.33 | 3303.40 |
| AT-VAC-4 | 35.54 | 793.42 | 34.77 | 1304.34 |
| AT-VAC-5 | 33.97 | 2178.44 | 33.21 | 3559.03 |
| AT-VAC-6 | 34.64 | 1421.23 | 33.01 | 4047.45 |
| Mean |  | 1539.61 ± 585.9 |  | 3223.92 ± 970.96 |

“-”, under the detection limit

### 3.2 Expression of specific immune-related genes

Different expression levels of three immunoglobulin genes encoding *IgM*, *IgD*, and *IgT* in the spleen and the head kidney were observed for the different treatment groups (Fig. 3). On day 14, a significant increase in *IgM* expression in the spleen (approximately 2.5 folds) was observed in the vaccinated groups that received pre-treatment with NB-O_3_ (NB-O_3_-VAC) compared with the unvaccinated groups (NB-O_3_-noVAC and AT-noVAC), while no significant difference was observed between any of the treatment groups on day 7 or day 21 post-vaccination. A significant upregulation in the spleen of *IgD* (approximately 2-fold) was found in the vaccinated groups that received pre-treatment with NB-O_3_ (NB-O_3_-VAC) at day 21 compared with the unvaccinated groups (NB-O_3_-noVAC and AT-noVAC), whereas no significant differences were observed in the spleen among all treatment groups at day 7 or 14 post-vaccination. On day 7 post-vaccination, neither the vaccinated nor the unvaccinated groups demonstrated substantial changes in *IgT* expression in the spleen of fish. *IgT* expression showed a higher relative change in both vaccinated groups compared to the unvaccinated groups at all time points. However, these differences were not statistically significant. At days 14 and 21 post-vaccination, *IgT* expression was substantially increased (approx. 2.8–4.1 folds) in the spleen of the NB-O_3_-VAC and AT-VAC groups compared to the fish in the NB-O_3_-noVAC and AT-noVAC groups. The highest expression was obtained in the NB-O_3_-VAC group (approximately 4.1-fold) at day 21 post-vaccination (Fig. 3A).

**Fig. 3.**
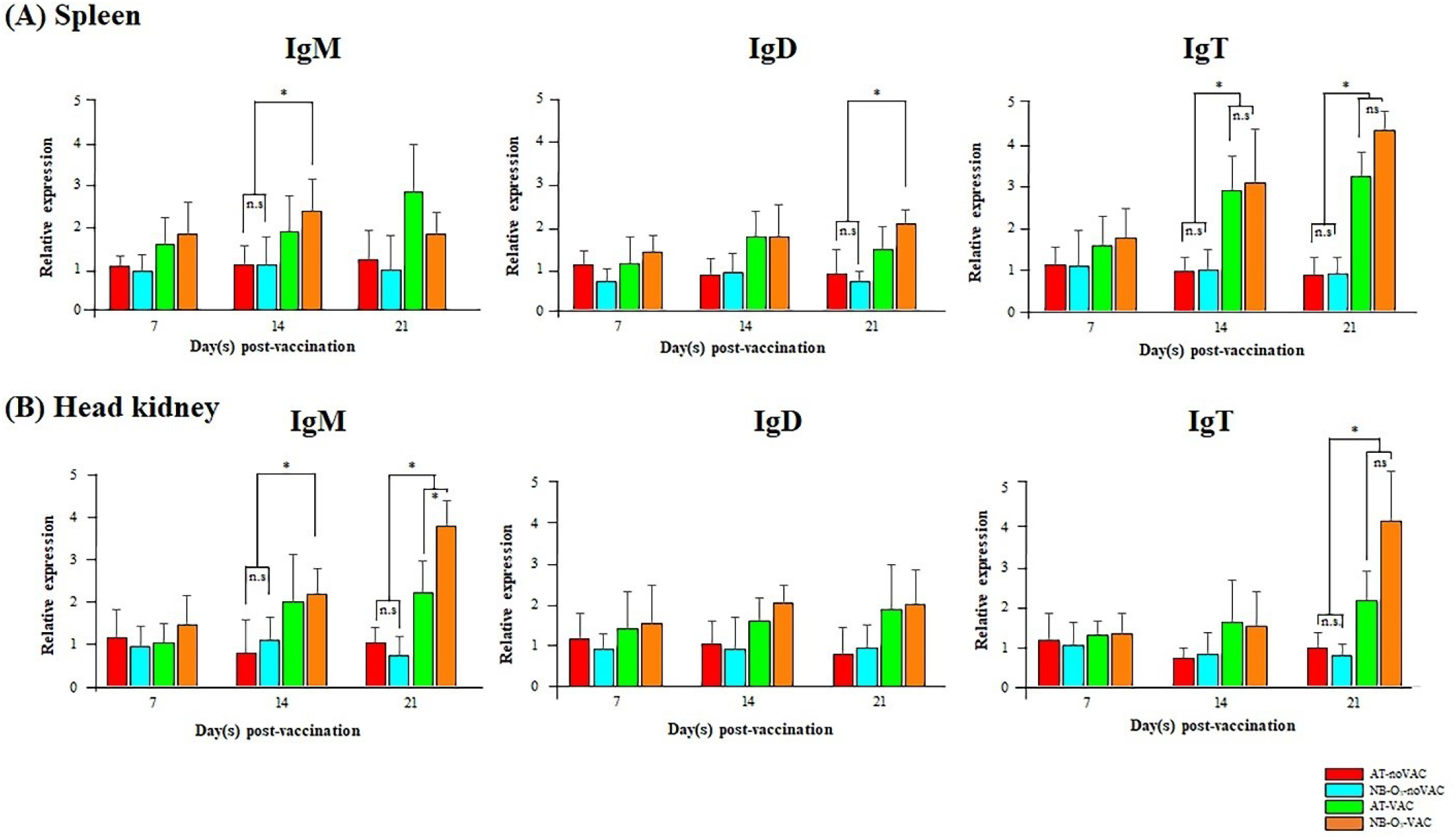
Comparative *IgM, IgD*, and *IgT* expression levels in the spleen and head kidney of the control and treated fish (n = 6) on days 7, 14, and 21 after immunization. The expression levels of three immunoglobulin genes were normalized by *β-actin*. Transcript levels of AT-noVAC groups at day 7 were set as 1. Error bar indicates standard deviation; “*” indicates a statistical significance (*P* < 0.05).

In the head kidney tissues, the relative expression of *IgM* was significantly increased in both the AT-VAC and NB-O_3_-VAC groups at day 14 (approx. 1.8–2.2 folds) and 21 (approx. 2.2–3.6 folds) post-vaccination, with significantly higher levels in the NB-O_3_-VAC fish. *IgM* expression in the NB-O_3_-VAC group was considerably upregulated compared with *IgM* expression in the AT-VAC group on day 21 post-vaccination. On day 7 post-vaccination, no significant variations in *IgM* expression were observed in any groups.

On days 7, 14, and 21 after immersion vaccination, *IgD* expression was essentially constant, and no alteration was detected in any of the treatment groups. There was a considerable increase in *IgT* of the AT-VAC and NB-O_3_-VAC groups (approx. 1.9–3.6 folds) compared with the NB-O_3_-noVAC and AT-noVAC groups at day 21 post-vaccination. Notably, expression levels of fish in the NB-O_3_-VAC groups were much higher than those of fish in the AT-VAC groups (Fig. 3B).

### 3.3 Analysis of specific-IgM antibody response

The systemic antibody response of the vaccinated fish showed substantially greater levels (*P* < 0.05) of specific IgM antibodies in their serum by indirect ELISA methods on days 7, 14, and 21 compared to fish that were not vaccinated. No significant specific antibody titer was detected in fish pre-vaccination (at day 0); however, the production of IgM serum in the vaccinated fish that received pre-treatment with NB-O_3_ was, on average slightly higher than that of vaccinated fish that did not receive pre-treatment with NB-O_3_ on day 7 (0.138 ± 0.005 vs. 0.116 ± 0.006), day 14 (0.144 ± 0.02 vs. 0.138 ± 0.015), and day 21 (0.146 ± 0.06 vs. 0.09 ± 0.005) post-vaccination, respectively. Similar to pre-vaccination fish, no significant systemic antibody response was detected in any of the unvaccinated fish on any of the sample days (Fig. 4A).

**Fig. 4.**
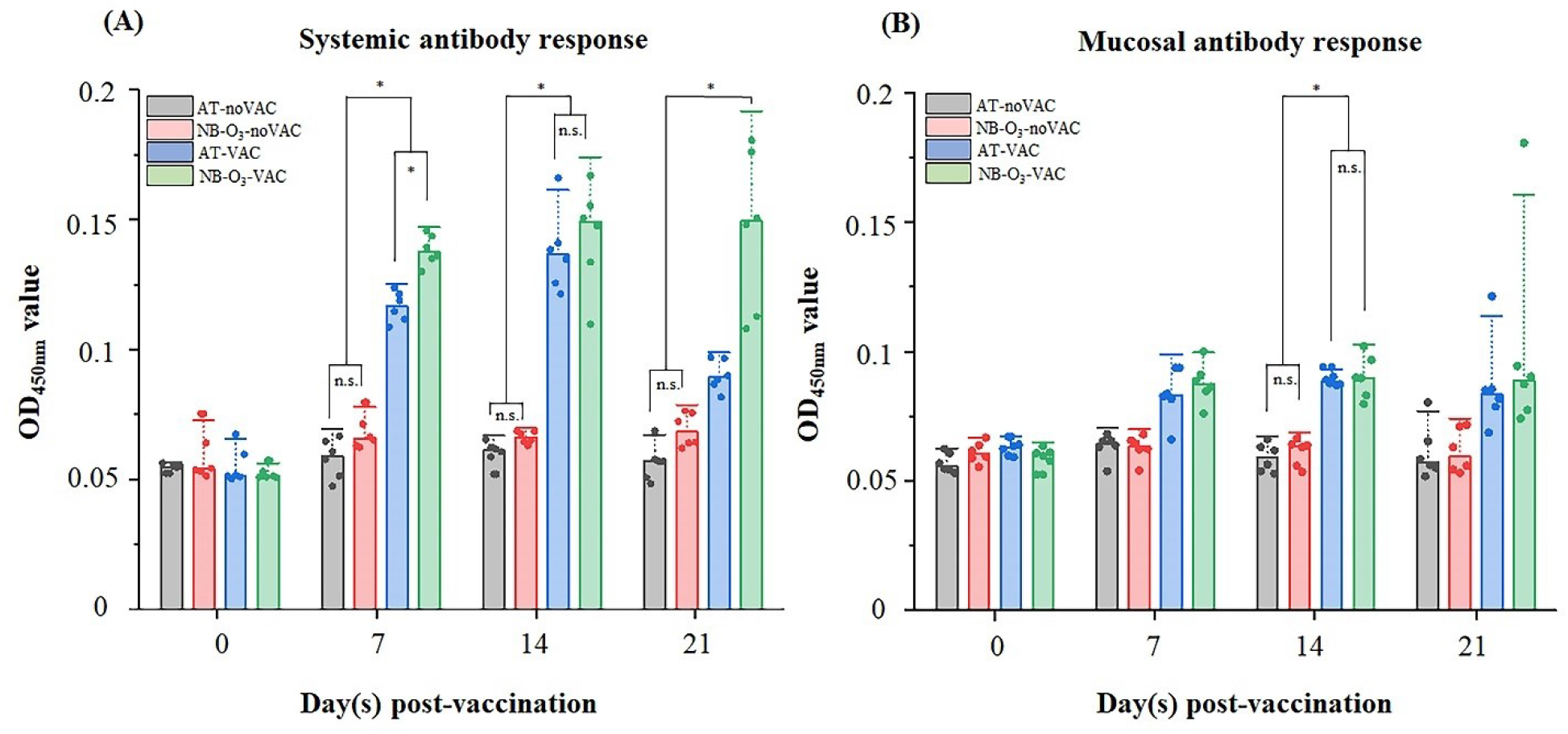
Specific antibody titer in the serum (A) and mucus (B) of Nile tilapia on days 0, 7, 14, and 21 of the immersion vaccination trials determined by ELISA. Serum and mucus antibody titers were determined using 1:512 and 1:16 dilutions, respectively. The optical density (OD) values were determined at 450 nm. Data are shown in mean ± standard deviation (SD), with each dot representing one biological replicate (n = 6). Statistical significance was determined by Kruskal– Wallis test. “*” indicates a statistical significance (*P* < 0.05), whereas “ns” indicates non-significant.

Analysis of IgM levels in the mucous membranes of tilapia after vaccination displayed a similar pattern to the systemic antibody response; however, lower OD_450 nm_ reading values of mucosal antibody titers were observed on days 7, 14, and 21 post-vaccination compared to the systemic antibody response measurements. Specific-IgM antibody responses in the vaccinated groups that received pre-treatment with NB-O_3_ were all slightly higher than those of the vaccinated group without pre-treatment with NB-O_3_ on days 7, 14, and 21. On day 14 post-vaccination, the mucosal IgM of fish in the vaccinated groups (NB-O_3_-VAC) had significantly higher antibody levels than those in the unvaccinated groups (NB-O_3_-noVAC and AT-noVAC). On the other days, all differences were not statistically significant (Fig. 4B).

### 3.4 Cumulative survival of vaccinated fish after challenging with *S. agalactiae*

Mortality was noted on day 3 post-challenge and persisted until day 6 and day 9 post-challenge for fish in the NB-O_3_-VAC and AT-VAC groups, respectively. However, fish mortality occurred earlier in the unvaccinated groups than in the vaccinated groups (on day 2) and persisted until day 8 post-challenge. The NB-O_3_-VAC group had the highest RPS of 70.5 ± 8.31%, followed by the RPS in the AT-VAC group (52.9 ± 16.63%) (Fig. 5A). This study demonstrated a significant difference (*P* = 0.003) between the NB-O_3_-VAC and the AT-noVAC groups. The AT-VAC groups also displayed a substantial difference compared to the AT-noVAC groups (*P* = 0.02) (Fig. 5B). Fish that died during the challenge experiment exhibited typical clinical signs of *S. agalactiae* infection. *Streptococcus agalactiae* was isolated from a representative group of moribund and dead fish on SSA.

**Fig. 5.**
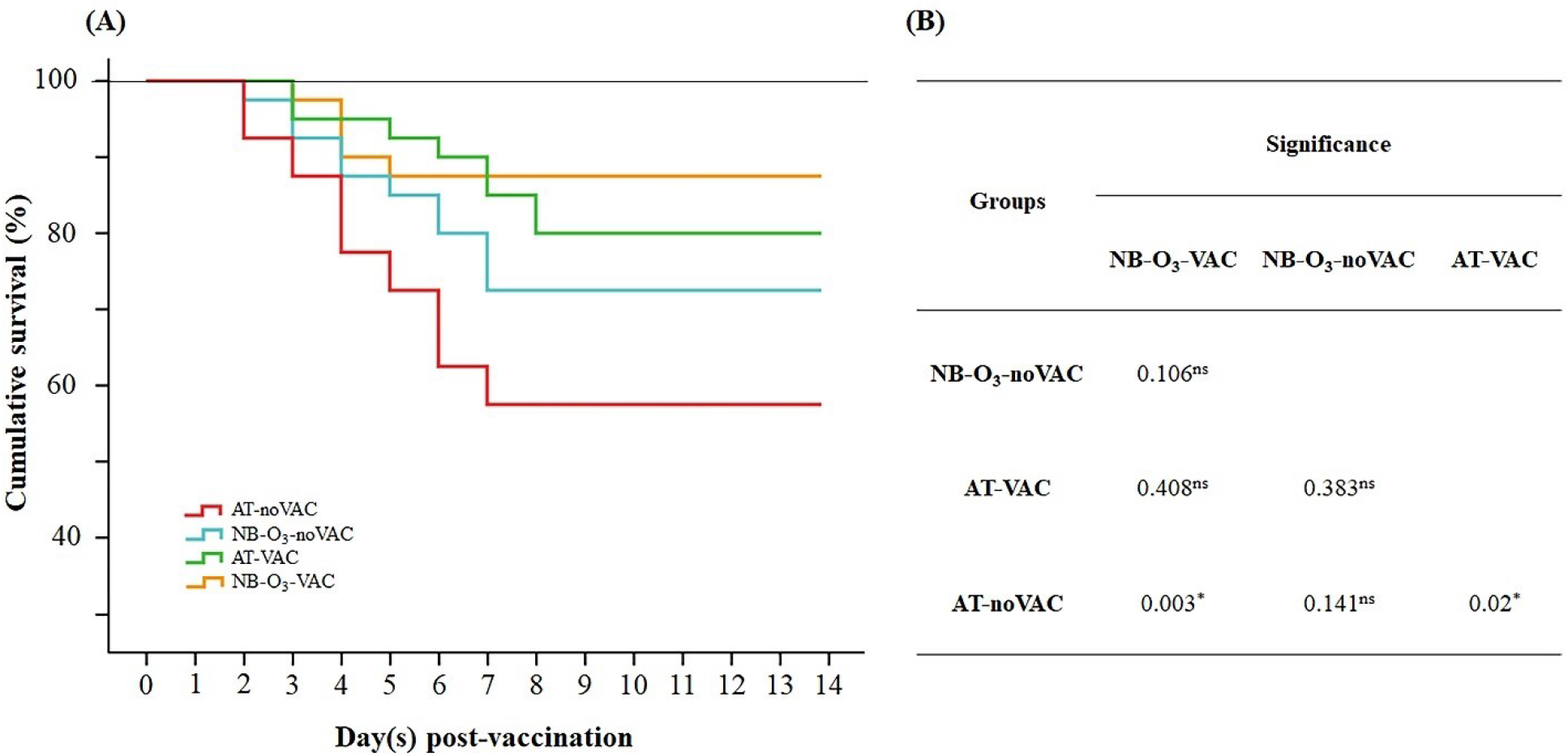
Kaplan–Meier analysis of (**A**) the cumulative survival of Nile tilapia (n = 40) challenged with *S. agalactiae* at day 21 post-vaccination. AT-noVAC, air-stone with no vaccine; NB-O_3_-noVAC, ozone nanobubbles without vaccine; AT-VAC, air-stone with vaccine; NB-O_3_-VAC, ozone nanobubbles with vaccine. The average cumulative survival of two trials is shown in the data. (**B**) The log-rank test was used to assess differences between the study treatments. “*” indicates a statistical significance (*P* < 0.05), whereas “ns” indicates non-significant.

## 4. Discussion

Based on the results of antigen uptake, antibody assay, expression levels of three immunoglobulin genes, and RPS, this study indicated that pre-treatment with NB-O_3_ improved immune responses in Nile tilapia to heat-killed *S. agalactiae* immersion vaccine. This approach represents a promising strategy for the prevention of streptococcus infections in Nile tilapia. Recently, several advantages of NB-O_3_ have been reported in terms of reducing the concentration of certain pathogenic bacteria and increasing dissolved oxygen (DO) in water. It has been suggested that NB-O_3_ could act as an “immunostimulant” to activate the fish innate immune system against bacterial infections [27] and improve the survivability of Nile tilapia challenged with *Aeromonas hydrophila* [25]. Notably, the findings of our investigation revealed that better bacterial antigen uptake into the gill tissues after 3 and 6 h immunization is achieved with fish in the vaccinated group that received a pre-treatment with NB-O_3_ (NB-O_3_-VAC) compared with fish in the AT-VAC group. It is possible that ozone nanobubbles in the NB-O_3_-VAC groups activated innate immune system, which modulated immune cells in response to infections [25–27]. When these innate immune cells are stimulated, they perform several response mechanisms to antigens, including phagocytosis, degranulation, and production of cytokines, which may trigger and/or recruit other leukocytes (e.g., neutrophils, macrophages, and dendritic cells) to the fish gills, and it is highly tempting to speculate that pre-treatment with NB-O_3_ might cause a temporary disruption of epithelial cells, leading to reducing barrier, increasing antigen uptake, and producing a greater immune response. Another possible explanation for the improved antigen absorption may be the physical and biological properties of ozone nanobubbles that require further investigation. Antigens may have been taken up by the gills or skins during immersion vaccination and processed by the innate immune system (e.g., phagocytes), where subsequent responses resulted in adaptive immune system and protected the immunized fish [34]. Indeed, pre-treatment with NB-O_3_ appears to have aided in the uptake of bacterial antigens during immersion vaccination. Several studies have previously demonstrated that enhanced antigen uptake results in increased protection [35,36]. These findings indicate that NB-O_3_ may be useful for improving the efficiency of immersion immunization in Nile tilapia.

In the current study, our data showed that fish in the vaccinated groups that received pre-treatment with NB-O_3_ had an increase in mRNA expression of three immunoglobulin classes (*IgM*, *IgD*, and *IgT*) compared to other treatment groups, including vaccinated fish that were not treated with NB-O_3_. Immunoglobulins (Ig) are important players in adaptive immune responses because they recognize and eliminate infections through a variety of mechanisms [37]. The immunoglobulin M (IgM) is the most abundant Ig in the plasma and the primary participant in systemic immunity, whereas the immunoglobulin T (IgT) is found in mucosal secretions and represented the primary Ig in mucosal immunity. The immunoglobulin D (IgD) is presumably involved in vertebrate immune responses, its relevance in Nile tilapia, however, remains unclear [38–40]. Specific immune gene expressions (*IgM*, *IgD*, and *IgT*) of fish in the vaccinated groups that received pre-treatment with NB-O_3_ were all slightly or significantly higher compared to fish in the other groups, reflecting the important role of NB-O_3_ in transporting vaccine antigens to the lymphoid organs and subsequent induction of mucosal and/or systemic immunity. Interestingly, our works demonstrate for the first time that immersion immunization induces a mucosal IgD response in Nile tilapia. The considerable elevations of *IgM*, *IgD*, and *IgT* in the spleens or head kidneys suggest that these immunoglobulins may play an essential role in defending fish against *S. agalactiae* infection.

The interaction between the cumulative survival of fish-immunized and antibody titers in the mucus and serum of Nile tilapia [41], Asian seabass [42], and Atlantic cod [43] highlights the critical role of antibody-mediated immunity in protecting fish against streptococcal infections. This concept was consistent with our observation; there were higher survival in the groups of fish with the higher antibody titers and these fish were more common in the NB-O_3_ treatment group [27]. The results obtained in this study are in agreement with a previous study, which induced significant protection in Asian seabass (*Lates calcarifer*), demonstrated by an RPS value ranging from 75–85% in both monovalent and bivalent vaccine groups after challenge with the inactivated *S. agalactiae* and *S. iniae* [42] or better RPS value (59.3 59.3% and 77.8%) was achieved with fish in the vaccinated group supplemented with adjuvants (aluminum hydroxide and FIA) compared with fish in the other groups without adjuvants [44]. Our previous study reported that pre-treatment with NB-O_3_ stimulated expression of immune-related genes of the fish innate immune system [27]. This may explain better responses of adaptive immune system after vaccination. Furthermore, the high RPS obtained in the current study in the NB-O_3_ group may be partially due to the contribution of specific immune mechanisms and the induction of mucosal and systemic IgM antibodies, as evidenced by upregulation of *IgM* in the spleens or head kidneys and an increase of total IgM in the mucus and serum of vaccinated fish.

There are still many questions to be addressed regarding the mechanisms of gene regulation as well as the mechanism of immunological responses in Nile tilapia. Further studies are needed to corroborate these findings and develop a deeper understanding of the mechanism of the immune stimulation observed in this study.

In conclusion, this study reported that pre-treatment with NB-O_3_ is a promising strategy for enhancing the efficacy of immersion vaccines against bacterial infections in tilapia, which has potential to be applied in aquaculture on a large scale.

## Data availability

The authors declare that they do not have any shared data available.

## Author contributions

**Nguyen Vu Linh**: Investigation, Methodology, Formal analysis, Writing – original draft, Software, and Resources. **Le Thanh Dien**: Investigation and Methodology. **Pattiya Sangpo**: Investigation and Methodology. **Saengchan Senapin**: Data curation and Writing - review & editing. **Anat Thapinta**: Supervision and Validation. **Wattana Panphut**: Supervision and Validation. **Sophie St-Hilaire**: Writing - review & editing, Funding acquisition, and Project administration. **Channarong Rodkhum**: Supervision and Validation. **Ha Thanh Dong**: Conceptualization, Data curation, Writing – review & editing, Supervision, Validation, Funding acquisition, and Project administration.

## Disclaimers

The views expressed herein do not necessarily represent those of IDRC or its Board of Governors.

## Acknowledgments

This research project received financial support from the UK government - Department of Health and Social Care (DHSC), Global AMR Innovation Fund (GAMRIF), and the International Development Research Center (IDRC), Ottawa, Canada. Nguyen Vu Linh has been supported by the Chulalongkorn University, Ratchadapisek Somphot Fund for Postdoctoral Fellowship. The authors like to express their gratitude to Mr. Nguyen Dinh-Hung for his technical support.

